# Computational design of CYP102A1 variants for the biosynthesis of a next-generation antiplatelet drug DT-678

**DOI:** 10.64898/2025.12.04.692440

**Authors:** Yudong Sun, Xiaoqiang Huang, Jifeng Zhang, Yoichi Osawa, Y. Eugene Chen, Haoming Zhang

## Abstract

Clopidogrel is a widely used antiplatelet prodrug to treat acute coronary syndromes. However, its clinical efficacy is hampered by ineffective bioactivation to produce the pharmacologically active metabolite (AM), leading to variability in antiplatelet response among different ethnic groups. To overcome the shortcomings of clopidogrel, DT-678 was developed by conjugating the AM to 3-nitropyridine-2-thiol via a mixed disulfide bond. It has been challenging to produce the conjugate in high yield by chemical synthesis. Here, we report the first de novo biosynthesis of DT-678 using engineered CYP102A1 variants. We applied structure-based computational design using UniDesign to generate three variants (UD4, UD5, and UD6) that enhanced the catalytic activity and selectivity toward DT-678 synthesis. Among them, UD6 demonstrated the highest total turnover number and DT-678-specific productivity under optimized conditions. Mechanistic analysis revealed that rapid enzyme inactivation, driven by reactive oxygen species (ROS) such as superoxide and hydrogen peroxide, limited overall yield. Remarkably, we found that ascorbic acid significantly protected CYP102A1 variants from inactivation and hence increased production yield. This work establishes a scalable enzymatic strategy for DT-678 biosynthesis and highlights the importance of combining protein engineering with redox control to overcome limitations in CYP-catalyzed reactions.

## Introduction

Clopidogrel is a widely used antiplatelet prodrug for the treatment of acute coronary syndromes ^1-2^. However, many patients experienced high on-treatment platelet reactivity, often due to genetic polymorphisms in cytochrome P450 (CYP) CYP2C19, a key enzyme responsible for converting clopidogrel to its pharmacologically active metabolite (AM) that irreversibly inhibits platelet P2Y^12^ receptor ^3-6^. Despite high bioavailability, only ∼15% of absorbed clopidogrel is metabolized by CYP enzymes, resulting in low levels of the AM estimated to be approximately 2% of ingested clopidogrel ^7^. As a result, individuals with CYP2C19 loss-of-function alleles and/or type 2 diabetes often respond poorly to clopidogrel ^4, 8-9^. Newer generation of antiplatelet drugs like prasugrel and ticagrelor are more efficacious but exhibit elevated risk of bleeding, underscoring the need for safer and more efficacious therapies ^10-13^.

To address this, we initially developed DT-678, a novel antiplatelet prodrug created by conjugating the AM to 3-nitropyridine-2-thiol (NPT). DT-678 contains two chiral centers (**Fig. 1**) and consists of a pair of stereoisomers (7S4R and 7R4S). This pair, referred to as PK2, can be separated by conventional high-performance liquid chromatography (HPLC) from another pair (7R4R and 7S4S) under Peak 1 (PK1) (**Fig. S1**). We formulated DT-678 using PK2 because of its higher potency in inhibiting platelets ^14-16^. The major advantage of DT-678 over clopidogrel is that it rapidly releases the AM by thiol exchange reaction with endogenous glutathione, bypassing CYP2C19-mediated activation ^17-18^. It achieves nearly complete AM conversion in all individuals and demonstrates superior efficacy with significantly lower bleeding risk in rabbits compared to existing antiplatelet drugs ^19^. In a Phase I clinical trial, DT-678 outperformed clopidogrel and is currently under development as a next-generation antiplatelet therapy ^20^. Due to the presence of two chiral centers, chemical synthesis of DT-678 remains challenging ^21-22^.

**Fig. 1.**
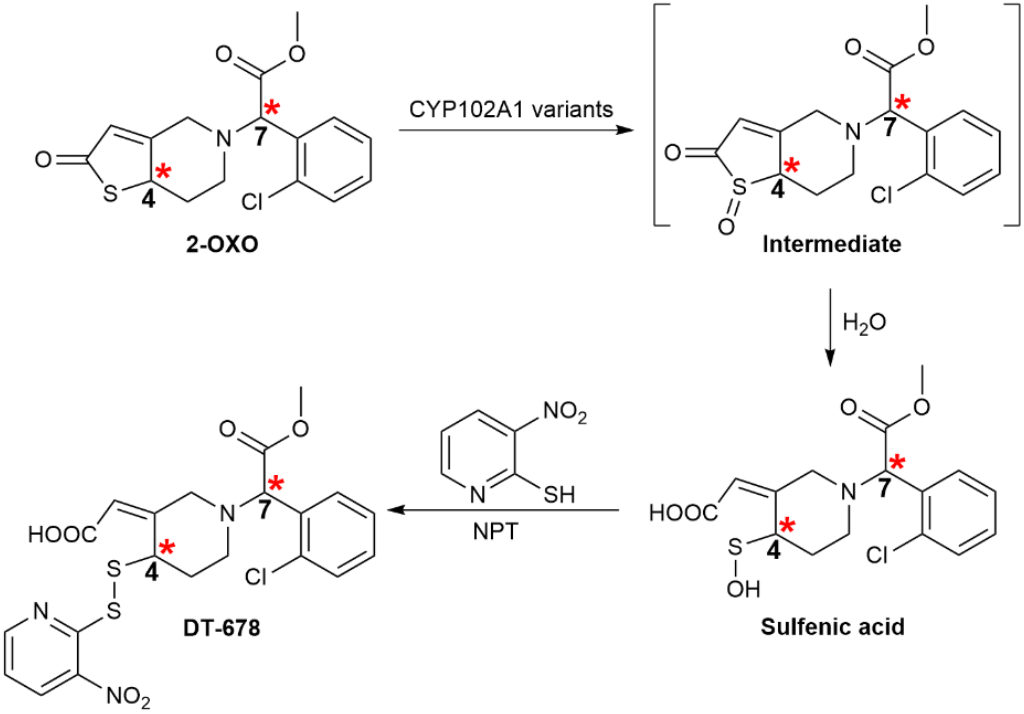
De novo biosynthesis of DT-678. The reaction starts with oxidation of racemic 2-OXO by CYP102A1 variants in the presence of NADPH and NPT at room temperature to form sulfenic acid intermediate which in turn conjugates with NPT to produce DT-678. As described in the main text, four stereoisomeric products are produced due to the presence of two chiral centers that are marked with red asterisks. DT-678 contains one pair of stereoisomers (7S4R, 7R4S).

In our recent studies, we developed UniDesign ^23-24^, a general protein design framework, and demonstrated its utility in redesigning CYP102A1 active-site architecture for highly stereoselective hydroxylation of omeprazole, another CYP2C19 substrate. Hereby we carried out de novo biosynthesis of DT-678 from commercially available 2-oxo-clopidogrel (2-OXO) using engineered CYP102A1 enzymes (**Fig. 1**). Based on a previously designed omeprazole-metabolizing triple variant (TM, A82F/F87V/L188Q), we used UniDesign to successively generate three variants: UD4 (TM + S72G), UD5 (UD4 + F87I) and UD6 (S72G/A82F/F87I). These variants show improved activity and selectivity for DT-678 synthesis. Furthermore, we demonstrate that ascorbic acid significantly extends the catalytic performance of CYP102A1 variants and increases the yield of DT-678 production.

## Results

### Designed CYP102A1 variants improve activity and selectivity for DT-678 synthesis

Previously we demonstrated that CYP102A1 variants A82F and TM metabolize omeprazole ^24^, exhibiting CYP2C19-like properties ^25^. Since 2-OXO is also metabolized by CYP2C19, we hypothesized that both variants would also catalyze the oxidation of 2-OXO. As shown in **Fig. 2**, the A82F variant achieved a total turnover number (TTN) of 7.8 (mol product per mol CYP enzyme) for production of all four isomers, with a DT-678-specific TTN (TTN_DT-678_) of 4.4, yielding a reaction selectivity of 1.4, defined as the PK2:PK1 ratio. In comparison, the TM variant considerably improved DT-678 synthesis with a TTN of 45.6 and a TTN_DT-678_ of 18.9, yielding a PK2:PK1 ratio of 0.7.

**Fig. 2.**
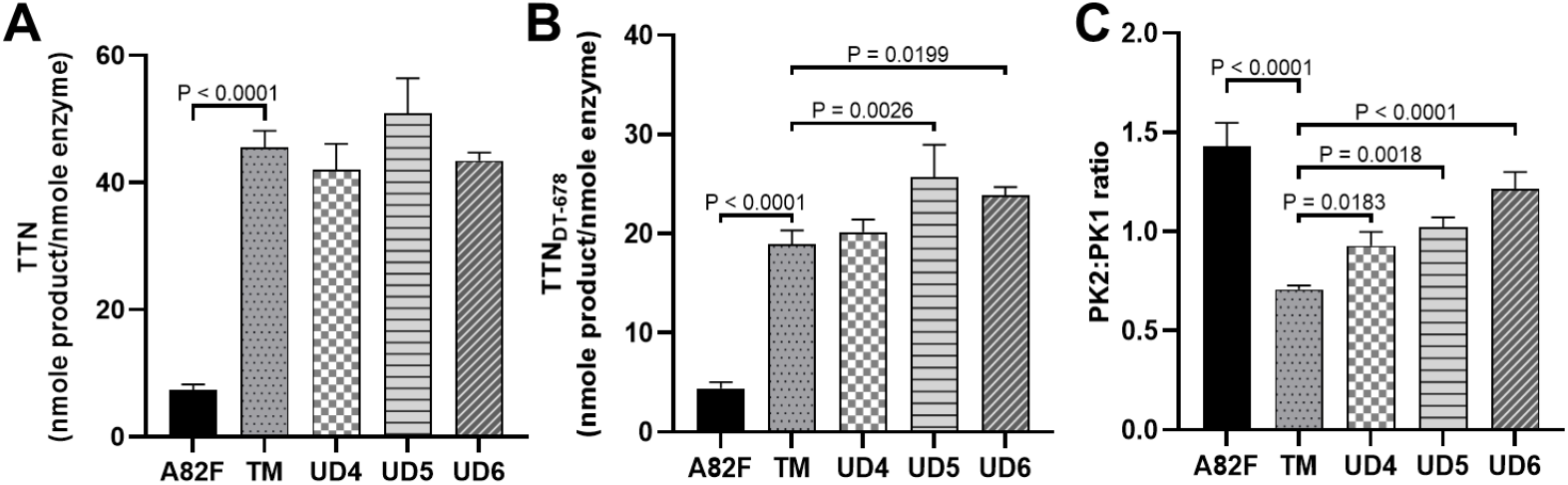
Productivity and selectivity of DT-678 biosynthesis by CYP102A1 variants. (A) Total turnover numbers (TTN) to produce the four isomers. (B) TTN specific to DT-678. (C) Stereoselectivity represented by the PK2:PK1 ratio. Product formation was quantified by HPLC in 2 h after initiation (n = 3). Statistical significance was assessed using ordinary one-way ANOVA in GraphPad Prism 10.

To improve the performance for DT-678 production, we used UniDesign to engineer the active-site residues of TM, following our previous strategy ^24^ (see Materials and Methods). Specifically, we aimed to introduce mutations favoring the binding of 7S4R over other isomers (7R4R, 7R4S, and 7S4S). UniDesign modeling predicted that introducing the S72G mutation into TM would result in a negative 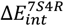 and positive 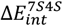 and 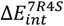, yielding strongly negative 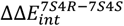 and 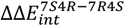 values (**Table 1**). This suggests that the new variant—UD4 (S72G/A82F/F87V/L188Q)—energetically favors the binding of 7S4R. Experimental validation showed that UD4 slightly reduced overall product formation but enhanced DT-678 production, with TTN and TTN_DT-678_ values of 42.0 and 20.1, respectively, and an improved PK2:PK1 ratio of 0.9 (**Fig. 2**). Similarly, UniDesign modeling predicted that replacing F87V with F87I in UD4 would further enhance the binding selectivity toward 7S4R by decreasing the interaction energies for 7S4S and 7R4S (**Table 1**). This led to the design of UD5 (S72G/A82F/F87I/L188Q). Experimentally, UD5 achieved TTN and TTN_DT-678_ of 50.9 and 25.7, respectively, and a PK2:PK1 ratio of 1.0, compared to 0.9 for UD4. Additionally, we generated another variant—UD6 (S72G/A82F/F87I)—by combining A82F with the S72G and F87I mutations, without direct UniDesign modeling. Interestingly, UD6 could achieve TTN and TTN_DT-678_ of 43.5 and 23.8, respectively. Although UD6 did not improve DT-678 production relative to UD5, it further increased the selectivity with a PK2:PK1 ratio of 1.2 (**Fig. 2**), likely by reducing activity toward 7S4S and/or 7R4R.

**Table 1.**
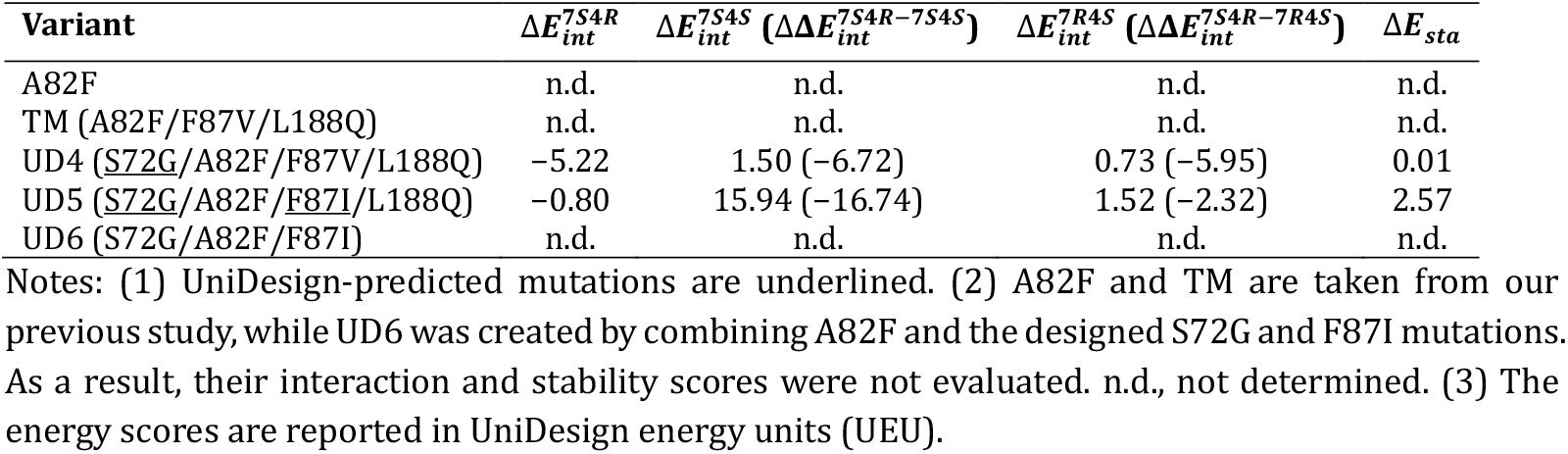
UniDesign-predicted stability and interaction energies of CYP102A1 variants.

| Variant | $\Delta E_{int}^{7S4R}$ | $\Delta E_{int}^{7S4S}$ ($\Delta\Delta E_{int}^{7S4R-7S4S}$) | $\Delta E_{int}^{7R4S}$ ($\Delta\Delta E_{int}^{7S4R-7R4S}$) | $\Delta E_{sta}$ |
| --- | --- | --- | --- | --- |
| A82F | n.d. | n.d. | n.d. | n.d. |
| TM (A82F/F87V/L188Q) | n.d. | n.d. | n.d. | n.d. |
| UD4 (<u>S72G</u>/A82F/F87V/L188Q) | -5.22 | 1.50 (-6.72) | 0.73 (-5.95) | 0.01 |
| UD5 (<u>S72G</u>/A82F/<u>F87I</u>/L188Q) | -0.80 | 15.94 (-16.74) | 1.52 (-2.32) | 2.57 |
| UD6 (S72G/A82F/F87I) | n.d. | n.d. | n.d. | n.d. |

Notes: (1) UniDesign-predicted mutations are underlined. (2) A82F and TM are taken from our previous study, while UD6 was created by combining A82F and the designed S72G and F87I mutations. As a result, their interaction and stability scores were not evaluated. n.d., not determined. (3) The energy scores are reported in UniDesign energy units (UEU).

### Reactive oxygen species (ROS) inactivates CYP102A1 variants

In our studies of the time course of the reaction, we found that the product formation by the UD6 variant plateaued in ∼40 min with a half-time of ∼7 minutes (**Fig. 3A**), despite the presence of excess NADPH. This observation suggested that CYP102A1 variants might be inhibited or inactivated over the course of the reaction. To investigate the cause for the loss of production, we measured the turnover rates for NADPH oxidation and ROS formation such as superoxide anion (O_2_^−^) and hydrogen peroxide (H_2_O_2_) (**Fig. 3B**). In the presence of 0.2 mM 2-OXO, the rates of NADPH oxidation and the O_2_^−^ and H_2_O_2_ formation were determined to be 1264.5 ± 51, 40 ± 0.01, and 46 ± 0.2 min^-1^, respectively. To assess whether ROS inactivated CYP102A1 variants, we determined the effect of superoxide dismutase (SOD) and catalase on the TTN for the UD6 variant. It is well established that SOD converts O_2_^−^ into H_2_O_2_ and O_2_, while catalase decomposes H_2_O_2_ into H_2_O and O_2_ ^26^. As shown in **Fig. 3C**, SOD substantially increased the TTN by approximately four folds. The combination of SOD and catalase exhibited an additive effect, further enhancing the TTN beyond SOD alone. Taken together, these results suggest that O_2_^−^ plays a predominant role in the inactivation of CYP102A1 variants.

**Fig. 3.**
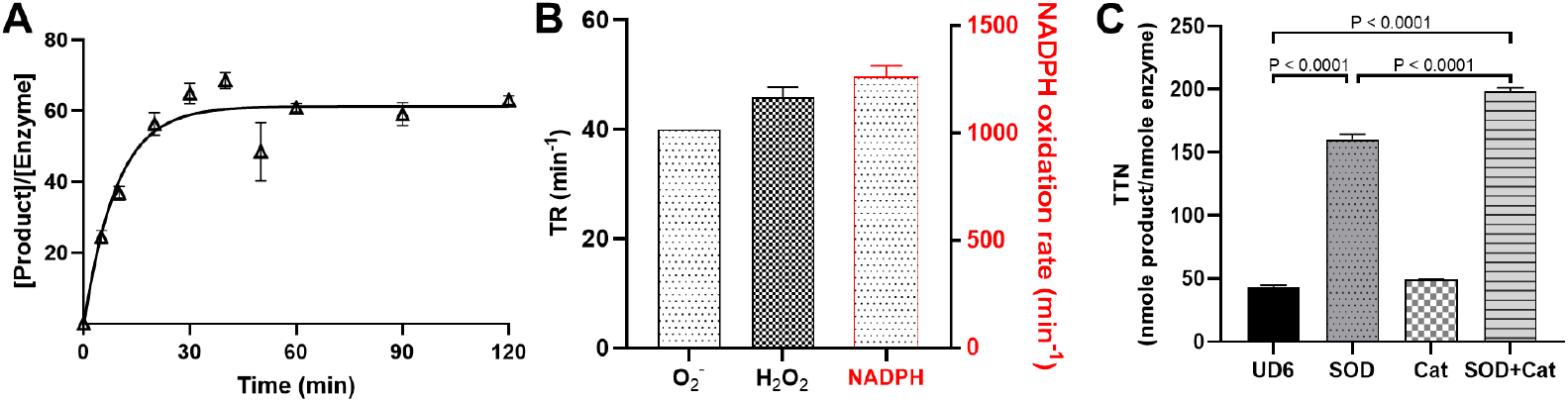
ROS inactivates CYP102A1 variants and reduces product formation. (A) Time-dependent formation of the four isomeric products by the UD6 variant. The reaction was carried out in the presence of 0.2 mM 2-OXO and the amount of the product was quantified by HPLC analysis as described in Materials and Methods. (B) The turnover rates for NADPH oxidation (marked in red, right Y-axis), and ROS formation (left Y-axis) for the UD6 variant. The rates were measured in the presence of 0.2 mM 2-OXO and calculated from the initial phase of the reaction in a linear range. (C) Effects of SOD and catalase (Cat) on the total turnover numbers (TTN) for the UD6 variant.

### Ascorbic acid protects CYP102A1 variants and significantly enhances DT-678 production

Ascorbic acid is a well-known antioxidant that neutralizes ROS such as O_2_^−^ and hydroxyl radical (•OH) ^26^. It also participates in redox reactions with H_2_O_2_, acting as a reducing agent to convert H_2_O_2_ into water. Based on these properties, we hypothesized that the addition of ascorbic acid could protect the CYP102A1 variants from oxidative inactivation and increase production yield.

To test this, we determined the TTN for the UD6 variant in presence of varying concentrations of ascorbic acid. The TTN achieved a peak value of 195.5 at 1 mM ascorbic acid and gradually decreased at elevated concentrations of ascorbate (**Fig. S2**). Thus, we determined the TTN and TTN_DT-678_ values, along with the PK2:PK1 ratios, for all CYP102A1 variants at 1 mM ascorbic acid (**Fig. 4**). As shown, each round of engineering led to progressive increases in the TTNs from 41.3 for A82F to 195.5 for UD6. A similar trend was observed in the improvement in stereoselectivity; the TM variant exhibited a PK2:PK1 ratio of 0.7, whereas the PK2:PK1 ratios of UD4, UD5, and UD6 were increased to 1.0, 1.2, and 1.3 respectively. The enhancements in both enzyme activity and selectivity contributed to significantly elevated TTN_DT-678_ values in the designed variants compared to TM. specifically, UD4 and UD5 achieved TTN_DT-678_ values of 77.2 and 95.8, respectively. UD6 was the most active variant, with a TTN_DT-678_ of 109.0, representing a 1.8-fold increase over TM and 4.4-fold increase over A82F.

**Fig. 4.**
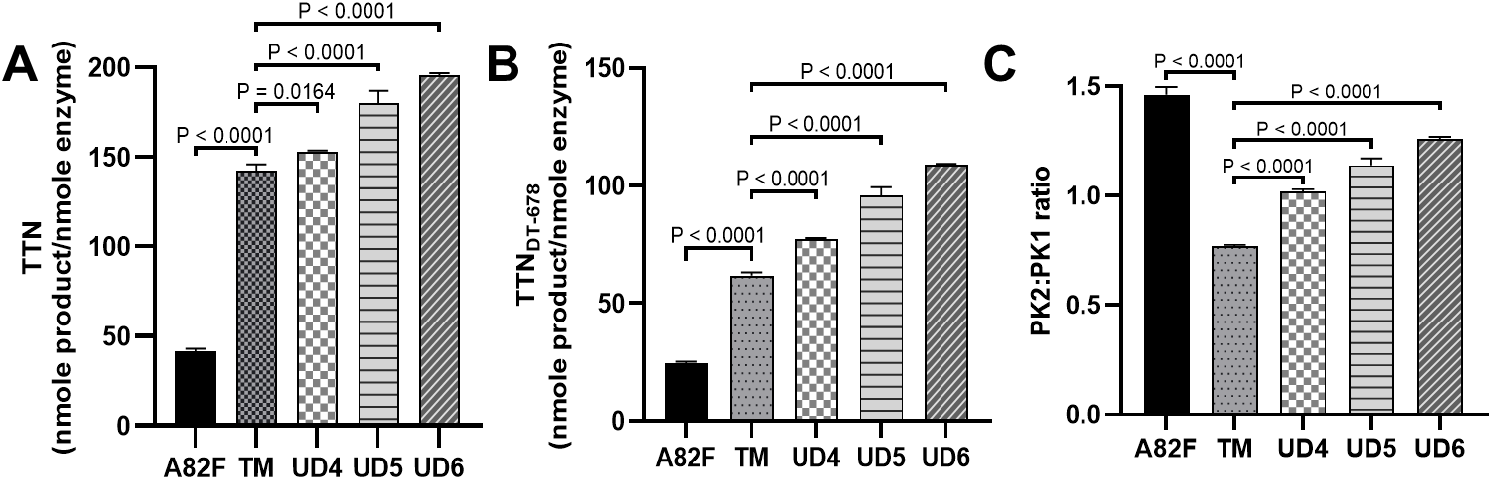
Productivity and selectivity of DT-678 synthesis by CYP102A1 variants in the presence of 1 mM ascorbate. (A) Total turnover numbers (TTN) for production of the four isomers. (B) TTN specific to DT-678. (C) Stereoselectivity represented by the PK2:PK1 ratio. Product formation was quantified by HPLC in 2 h after initiation (n = 3). Statistical significance was assessed using ordinary one-way ANOVA in GraphPad Prism 10.

### Designed CYP102A1 variants improve catalytic efficiency for 2-OXO oxidation

To better understand the improved production of DT-678 by the designed variants, we determined the kinetic parameters for 2-OXO oxidation in the presence of 1 mM ascorbic acid. The results are summarized in **Table 2**. Despite being a phenotypical CYP2C19 variant, A82F shows a moderate *k*_cat_ of 5.0 ± 0.05 min^−1^ with a catalytic efficiency of 0.04 min^−1^·μM^−1^. The TM variant, which served as the template variant for UniDesign, shows a 3.5-fold increase in *k*_cat_ and 2.5-fold improvement in catalytic efficiency compared to A82F. With each round of engineering, the *k*_cat_ values of the designed variants progressively increased; notably, the UD6 variant achieved a nearly twofold increase in *k*_cat_ to 33.6 ± 0.6 min^−1^, along with a 40% enhancement in catalytic efficiency. However, these improvements come with a tradeoff: two rounds of engineering resulted in ∼8–38% increase in *K*_M_, partially offsetting the gains in catalytic efficiency.

**Table 2.** Kinetic parameters for 2-OXO oxidation to form all four isomers by CYP102A1 variants in presence of 1 mM ascorbic acid. Products were analyzed by HPLC as described in Materials and Methos.

| Variant | $k_{cat}$ (min<sup>-1</sup>) | $K_M$ (μM) | $k_{cat}/K_M$ (min<sup>-1</sup>μM<sup>-1</sup>) |
| --- | --- | --- | --- |
| A82F | 5.00 ± 0.05 | 121.3 ± 6.11 | 0.04 |
| TM (A82F/F87V/L88Q) | 17.6 ± 2.3 | 174.0 ± 23.3 | 0.10 |
| UD4 (S72G/A82F/F87V/L88Q) | 20.7 ± 2.2 | 187.9 ± 39.4 | 0.11 |
| UD5 (S72G/A82F/F87I/L88Q) | 27.2 ± 0.9 | 192.2 ± 13.8 | 0.14 |
| UD6 (S72G/A82F/F87I) | 33.6 ± 0.6 | 241.9 ± 1.2 | 0.14 |

## Discussion

In this study, we report the de novo biosynthesis of DT-678, a next-generation antiplatelet drug, from 2-OXO, a commercially available intermediate. This pathway utilizes engineered CYP102A1 enzymes to mimic CYP2C19-mediated clopidogrel bioactivation in vivo. To enable practical DT-678 production, we characterized the engineered CYP102A1 variants and optimized the reaction conditions to enhance their catalytic stability. The production yield has been substantially increased by the designed variants in the presence of ascorbic acid.

The TM variant was originally developed in our previous study for the hydroxylation of omeprazole ^24^. Interestingly, we found that TM also exhibited significantly higher activity toward 2-OXO metabolism compared to the A82F variant. Given that both omeprazole and 2-OXO are known substrates of CYP2C19, we hypothesize that TM may also catalyze the oxidation of other CYP2C19 substrates and serve as a broadly useful scaffold for further engineering toward alternative substrates.

Although the TM variant shows promising activity in converting 2-OXO to DT-678, its performance remains suboptimal. To improve both activity and selectivity, we applied iterative computational design using UniDesign and identified two key mutations at residues 72 and 87. These mutations led to the development of the UD4, UD5, and UD6 variants, which showed progressively enhanced activity and selectivity for DT-678 synthesis. Compared to the baseline A82F and starting TM variants, these results highlight how UniDesign-enabled active-site engineering can effectively reshape enzyme– substrate interactions and catalytic outcomes, consistent with our previous study ^24^.

Residue 72 is located near the entrance of the substrate access channel, while residue 87 resides within the active-site pocket (**Fig. 5A**). The S72G mutation likely enlarges the tunnel and permits a displacement of the 7S4R isomer (**Fig. 5B**), allowing it to adopt a conformation more favorable for catalysis. This structural adjustment appears specific to 7S4R, as the binding poses of 7S4S (**Fig. 5C**) or 7R4S (**Fig. 5D**) remain largely unchanged. Notably, S72G also disrupts a favorable hydrogen bond between the native Ser72 and 7S4S, as well as a weaker interaction with 7R4S, potentially further disfavoring these isomers. Structural superposition of the binding models suggests that Val87 in the enzyme–7S4R complex sterically clashes with the 7S4S pose and is near 7R4S (**Fig. 5E**), indicating that a bulkier residue at position 87 may further disfavor 7S4S and 7R4S.

**Fig. 5.**
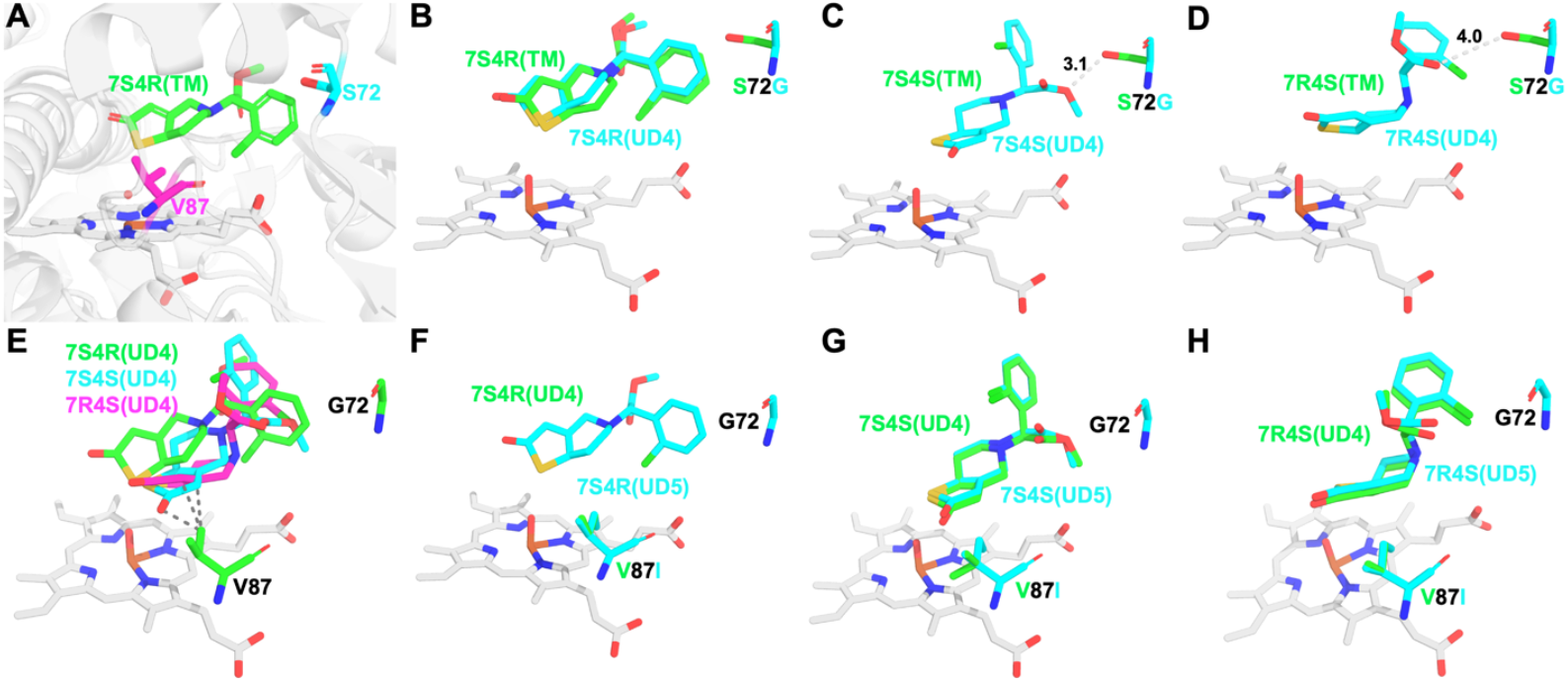
UniDesign modeling of CYP102A1 variants. (A) Structural locations of residue 72 and 87 in the enzyme. (B) Comparison of 7S4R binding poses between the TM and UD4 variants. (C) Comparison of 7S4S binding poses in TM and UD4; in TM, Ser72 forms a favorable hydrogen bond with 7S4S. (D) Comparison of 7R4S binding poses in TM and UD4; in TM, Ser72 forms a weak hydrogen bond with 7R4S. (E) Superposition of 7S4R, 7S4S, and 7R4S poses in the UD4 variant, highlighting spatial differences. The V87 in the 7S4R model forms clashes with the 7S4S and 7R4S poses. (F) Comparison of 7S4R binding poses between UD4 and UD5. (G) Comparison of 7S4S binding poses in UD4 and UD5. (H) Comparison of 7R4S binding poses in UD4 and UD5.

Consistent with this prediction, replacing UD4’s Val87 with Ile in UD5 did not affect the binding pose of 7S4R relative to UD4 (**Fig. 5F**), but caused noticeable displacement of both 7R4S (**Fig. 5G**) and 7S4S (**Fig. 5H**), thereby decreasing their binding favorability. This suggests that the improved selectivity observed in UD5 and UD6 stems primarily from disfavoring these undesired isomers. Interestingly, UniDesign’s suggestion to introduce a bulkier residue at position 87 aligns with the high selectivity observed in the A82F variant, which retains the even bulkier native Phe87. However, the presence of Phe87 leads to reduced catalytic activity, indicating an tradeoff between enzyme activity and selectivity in DT-678 synthesis.

The effects of successive engineering are manifested in the kinetic parameters of these variants (**Table 2**). The baseline variant A82F exhibits the lowest *k*_cat_ of 5 min^-1^ and a catalytic efficiency of 0.04 min^-1^·µM^-1^, resulting in poor TTN for DT-678 production (**Fig. 4C**). In contrast, the TM variant, taken from our prior work, shows elevated *k*_cat_ and catalytic efficiency, affording a good starting point for further engineering. Successfully, each round of engineering progressively enhances the *k*_cat_ and catalytic efficiency. Ultimately the UD6 variant shows a 1.9-fold and 6.7-fold increase in *k*_cat_ over the TM and A82F variants, respectively.

The inactivation of CYP102A1 variants appears to be a major bottleneck in DT-678 production, as we observed a complete loss of enzyme activity within approximately 40 min of the reaction (**Fig. 3A**), significantly limiting overall yield. To explore the underlying cause, we monitored ROS levels during the reaction and found substantial accumulation of both O_2_^−^ and H_2_O_2_. Notably, the addition of superoxide dismutase (SOD), but not catalase, markedly enhanced DT-678 production (**Fig. 3C**), suggesting that O_2_^−^ produced in the oxidative uncoupling process plays a key role in CYP enzyme inactivation ^27^. The detrimental effects of ROS can be mitigated by ascorbic acid, which likely protects the enzyme by scavenging ROS and preserving its structural integrity ^28^. These findings underscore the critical importance of protecting CYP102A1 variants from oxidative damage in practical biocatalytic applications.

In conclusion, we established the first biosynthetic route for DT-678 production using engineered CYP102A1 variants, enabling direct conversion from 2-OXO with improved activity and selectivity. Through structure-guided design with UniDesign, key mutations were introduced to enhance stereoselectivity toward the active 7S4R isomer. Additionally, we identified oxidative inactivation, primarily driven by O_2_^−^, as a major limitation to enzyme performance, which was effectively mitigated by ascorbic acid. These findings provide a foundation for scalable, enzyme-based DT-678 synthesis and demonstrate the potential of combining protein engineering with redox management to optimize P450 biocatalysis.

## Material and Methods

### Chemicals

All chemicals used are of the highest purity available. NADPH, NADP^+^, ascorbic acid, and D-glucose were purchased from Sigma-Aldrich (St Louis, MO). Racemic 2-OXO was from Toronto Research Company (Ontario, Canada). Carbon monoxide gas was from Cryogenic Gas (Detroit, MI).

### Computational design of CYP102A1 variants

We aimed to redesign the CYP102A1 TM variant to enhance its catalytic activity and selectivity for DT-678 synthesis based on the crystal structure of the CYP102A1 heme domain (PDB ID: 5XA3). Given the well-established reaction mechanism (**Fig. 1**), we modeled the interactions of CYP102A1 with all four isomers - 7S4R, 7S4S, 7R4S, and 7R4R. Because of the high structural similarity between the four stereoisomers and 2-OXO, we generated the four isomers by modifying the 2-OXO structure (PubChem {Kim, 2013 #29} ID: 56848893), followed by energy minimization and atomic charge assignment using Chimera ^29^, as described in our previous work ^24^.

Using the 5XA3 structure as a scaffold, we generated ligand poses for each isomer via a grow-and-check approach ^24^. This process yielded 3132, 317, 752, and 0 viable poses for 7S4R, 7S4S, 7R4S, and 7R4R, respectively, under predefined energy thresholds (100 for “Internal” and 15 for “Backbone” energies). The absence of valid poses for 7R4R indicated that it was not physically accommodated within the 5XA3 scaffold and was therefore excluded from subsequent mutational design analyses. In the first round of design, based on the TM variant, we selected 19 residues (positions 26, 72, 74, 75, 78, 82, 87, 181, 185, 188, 263, 264, 328, 329, 330, 332, 354, 437, and 438) for mutation, along with 14 additional sites (positions 25, 70, 71, 88, 177, 260, 266, 267, 355, 356, 357, 435, 436, and 439) for side-chain repacking. Each of the 19 mutable sites was substituted with all 19 alternative amino acids, resulting in a total of 361 mutants.

For each variant, we used UniDesign to repack the enzyme–isomer complex and minimized the system’s total energy via simulated annealing Monte Carlo (SAMC) simulations ^30^. Ligand poses were sampled from the previously generated ensembles (7S4R, 7S4S, and 7R4S), and amino-acid side-chain conformations were taken from the Dunbrack2010 backbone-dependent rotamer library ^31^. The final ligand pose was determined jointly with protein side-chain rotamers, and both total system energy and enzyme–ligand interaction energy were calculated. For each mutant, three independent SAMC trajectories were performed, and the design with the lowest total energy was selected. To triage these variants, we focused on those that improved the interaction energy (*E*_*int*_) for the preferred 7S4R isomer—identified as the most potent—without significantly compromising enzyme structural stability (*E*_*sta*_) relative to the reference TM variant. The selection criteria were as follows:

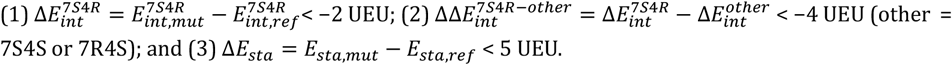

In the second round, based on the UD4 variant, we fixed positions 72 and 188 to Gly and Gln, respectively, generating 323 mutants. The selection criteria were slightly relaxed to reVlect the improved reference background: (1) 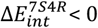 UEU; (2) 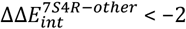 UEU; and (3) Δ*E*_*sta*_ < 5 UEU.

### Construction, expression, and purification of CYP102A1 variants

Target mutations were introduced into template plasmid using Quikchange site-directed mutagenesis kit, following the manufacturer’s instruction (Agilent Technologies). We constructed the UD4 mutant by introducing the S72G mutation into the plasmid cDNA of pCWori-CYP102A1-TM-6×His that we prepared previously ^24^, in which a 6× His tag was inserted after the starting codon ATG to facilitate purification using immobilized metal affinity chromatography. Then, we introduced the V87I mutation into the plasmid cDNA of pCWori-CYP102A1-UD4-6×His to construct UD5. Later, the F87I and S72G mutations were introduced into the plasmid cDNA of pCWori-CYP102A1-A82F-6×His to construct UD6. A list of site-directed mutagenic primers used to construct these mutants is provided in Table S1. The constructed variants were over-expressed in *Escherichia coli* C41(DE3) cells and purified on a PROTEINDEX Ni-Penta™ Agarose affinity column (Marvelgent Biosciences, Waltham, MA), as previously described ^24^. The concentration of CYP102A1 variants were determined by the method of Omura and Sato ^32^.

### Measurement of the rate of NADPH oxidation

The rate of NADPH oxidation was determined as previously described ^24^. CYP102A1 variants (0.05 μM) and varying concentrations of racemic 2-OXO (0–0.3 mM) were pre-equilibrated in 0.5 mL of 0.1 M potassium phosphate (KPi) buffer, pH 7.4, for 5 min at 25 °C. The reactions were initiated by the addition of 0.2 mM NADPH and monitored at 340 nm for 60 s on a UV-visible spectrophotometer (Shimadzu PC 2043 UV). The kinetic parameters *K*_M_ and *k*_cat_ were obtained by fitting the initial velocities to the Michaelis–Menten equation using GraphPad Prism 10 at various 2-OXO concentrations.

### Measurement of the turnover rate and total turnover number (TTN) in DT-678 production

The total turnover numbers were determined in 0.1 mL of 0.1 M KPi buffer containing 0.5 μM CYP102A1 variants, 0.2 mM of 2-OXO, 0.3 mM NPT, 0.5 mM NADP^+^, 10 mM glucose, and varying concentrations of ascorbic acid (when present). After thermal equilibration at 25 °C for 5 min, aliquots of glucose dehydrogenase were added at 100 μg/mL to initiate the reaction. The reaction was then terminated in 3 h with 0.1 mL acetonitrile. All reactions were performed in triplicates.

After centrifuged at 20,000×*g* for 20 min, the samples were analyzed by HPLC to quantify the amount of product DT-678 on a Shimadzu binary HPLC system equipped with a photodiode array detector and autosampler. Product DT-678 was separated from reactant 2-OXO on a reverse phase C18 column (Zorbax C18, 3×150 mm, Agilent Technologies) at a flow rate of 0.4 ml/min in a binary mobile phase consisting of 0.1 % formic acid in water (A) and 80% acetonitrile:20% methanol (B). The gradient was as follows: 30% B for 2 min, increasing to 40% B in 8 min, 75% in 5 min, 90% in 1 min and held at 90% for 5 min. The C18 column was maintained at 37 °C. The identity of product DT-678 was confirmed by comparison of its MS^2^ spectrum with authentic DT-678 standard (**Fig. S3**).

### Measurement of the rate of H_2_O_2_ production

The primary reaction mixture consists of 0.05 μM CYP102A1 variants, 0.2 mM 2-OXO and 0.3 mM NPT in 0.1 M KPi buffer (pH 7.4). The reactions were initiated by the addition of 0.2 mM NADPH and then quenched with 5 μL of 3.2 M hydrochloric acid in 2 min. The amount of H_2_O_2_ produced in the reaction was then quantified using ferric-xylenol orange colorimetric assay as previously described ^33^. The assay solution consists of 0.125 mM xylenol orange, 25 mM sulfuric acid and 200 mM ferrous ammonium acetate. Aliquots of 40 μL quenched primary reaction mixture were immediately transferred to 160 μL assay solution and the absorbance at 560 nm were recorded. Quantitation of H_2_O_2_ was based on calibration with known amount of H_2_O_2_ standards.

### Measurement of the rate of O_2_^−^ production

Succinylated cyt c was used to quantify the amount of superoxide as previously reported ^34-35^. In a typical reaction, CYP102A1 variants (0.05 μM), rat P450 reductase (0.01 μM, when present), various concentrations of racemic 2-OXO (0–0.3 mM), and 25 μM naïve or succinylated cyt c were pre-equilibrated at 25°C for 5 minutes in 0.5 mL of 0.1M KPi buffer (pH 7.4). The reaction was then initiated by the addition of 0.2 mM NADPH. The absorbance change was monitored for 1 min at 550 nm for the reduction of succinylated cyt c by superoxide. An extinction coefficient of 21 mM^-1^cm^-1^ for reduced cyt c was used to quantitate the amount of superoxide produced in the reaction.

## Data availability

All data are available upon request to the corresponding authors.

## Acknowledgments

This work was in part supported by grants from the National Institutes of Health (GM149016 to X.H.; HL159900 and HL159871 to Y.E.C.; GM153714 to Y.O., and ES030791 to H.Z.). H.Z. acknowledged the support by an intramural grant from the University of Michigan Medical School. The computational research in this work was supported through the computational resources and services provided by Advanced Research Computing at the University of Michigan.

## Author contributions

X.H., Y.E.C., and H.Z. conceived and administrated the project. Y.E.C. and H.Z. supervised the project. X.H. developed UniDesign and conducted computational enzyme design. Y.S. performed molecular experiments and biochemical assays. Y.S. and X.H. wrote the initial manuscript. X.H., J.Z., Y.O., Y.E.C., and H.Z. edited the manuscript. All authors discussed the results and contributed to the final manuscript.

## Supporting Information

**Fig. S1.**
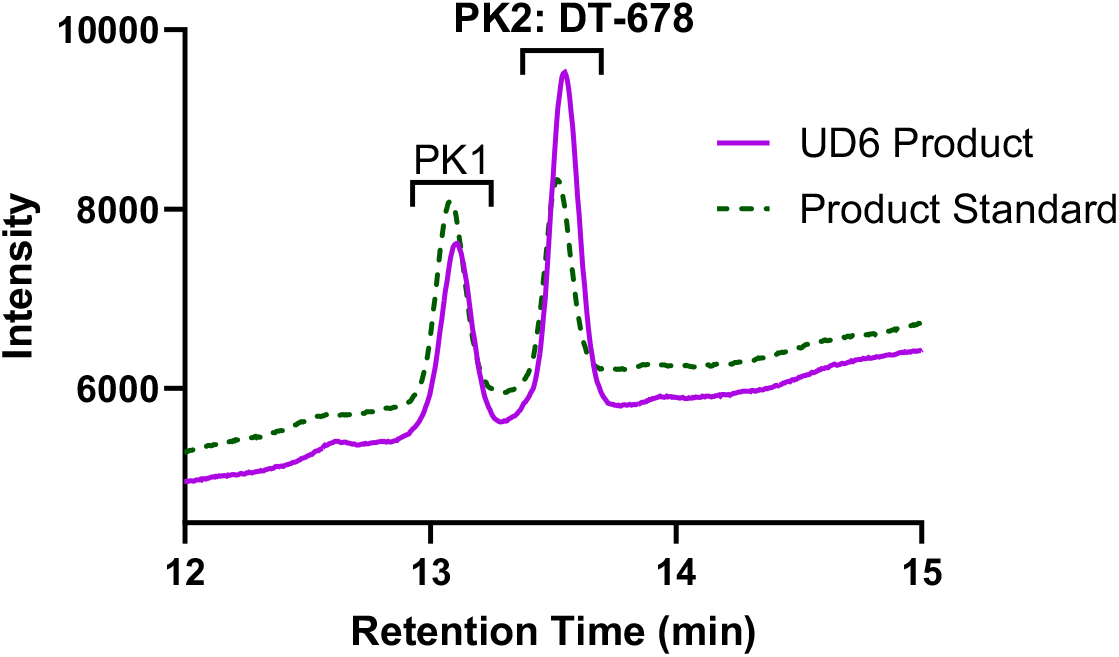
Representative HPLC elution profiles of the reaction product and product standard. The reaction product by the UD6 variant (solid purple trace) and product standard (dashed green trace) are resolved into two distinct peaks, each of which represents a pair of isomers: PK2 contains the 7S4R and 7R4S isomers in DT-678 and PK1 contains the 7S4S and 7R4R isomers. Product standards were generous gifts from Beijing SL Pharmaceuticals (Beijing, China).

**Fig. S2.**
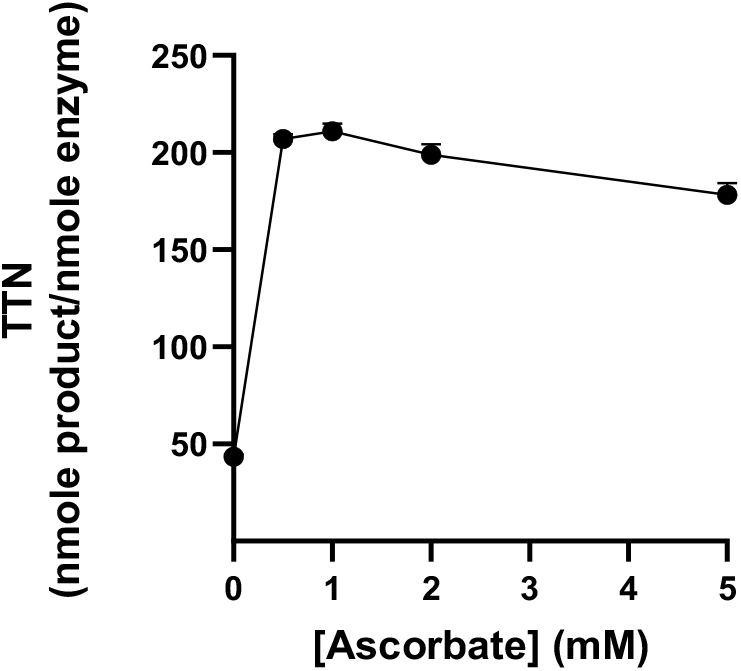
Dependence of the total turnover numbers (TTN) of the UD6 variant on the concentration of ascorbate. The TTN was determined at 25 °C in the presence of 0.2 mM 2-OXO as described in Materials and methods.

**Fig. S3.**
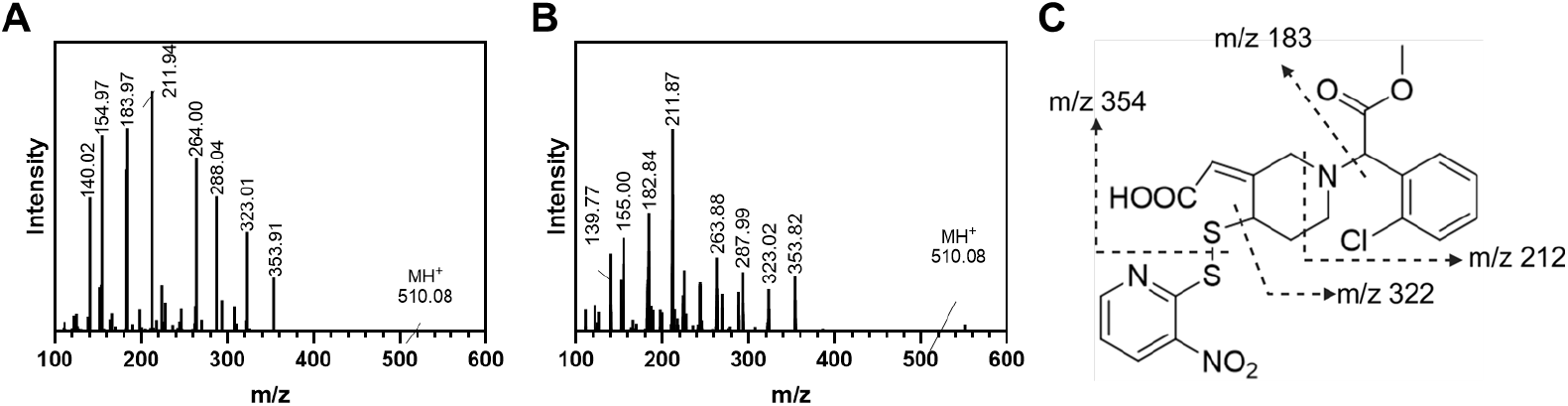
MS^2^ spectra of DT-678 produced by the UD6 variant (A) and DT-678 standard (B), with corresponding fragmentation pattern (C). The MS^2^ spectra was recorded with the following parameters: spray voltage of 3.5 kV, vapor temperature of 300 °C, capillary temperature of 290 °C, and collision energy of 35%.

**Table S1.**
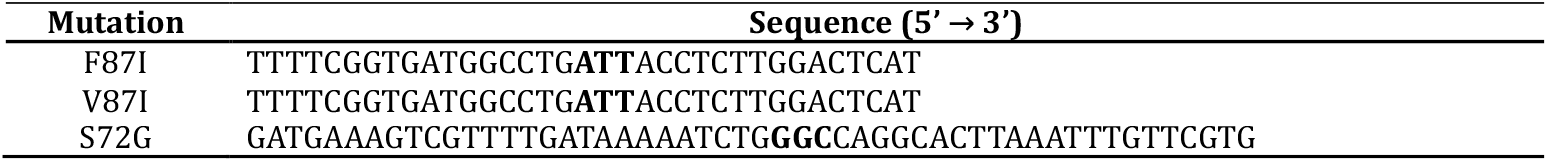
A list of site-directed mutagenic primers used to construct UD4, UD5 and UD6 mutants. The codons in bold are mismatches that indicate designated mutations.

## References

1. Rao, S. V.; O’Donoghue, M. L.; Ruel, M.; Rab, T.; Tamis-Holland, J. E.; Alexander, J. H.; Baber, U.; Baker, H.; Cohen, M. G.; Cruz-Ruiz, M.; Davis, L. L.; de Lemos, J. A.; DeWald, T. A.; Elgendy, I. Y.; Feldman, D. N.; Goyal, A.; Isiadinso, I.; Menon, V.; Morrow, D. A.; Mukherjee, D.; Platz, E.; Promes, S. B.; Sandner, S.; Sandoval, Y.; Schunder, R.; Shah, B.; Stopyra, J. P.; Talbot, A. W.; Taub, P. R.; Williams, M. S., 2025 ACC/AHA/ACEP/NAEMSP/SCAI Guideline for the Management of Patients With Acute Coronary Syndromes: A Report of the American College of Cardiology/American Heart Association Joint Committee on Clinical Practice Guidelines. Circulation 2025, 151 (13), e771–e862.

2. Byrne, R. A.; Rossello, X.; Coughlan, J. J.; Barbato, E.; Berry, C.; Chieffo, A.; Claeys, M. J.; Dan, G. A.; Dweck, M. R.; Galbraith, M.; Gilard, M.; Hinterbuchner, L.; Jankowska, E. A.; Juni, P.; Kimura, T.; Kunadian, V.; Leosdottir, M.; Lorusso, R.; Pedretti, R. F. E.; Rigopoulos, A. G.; Rubini Gimenez, M.; Thiele, H.; Vranckx, P.; Wassmann, S.; Wenger, N. K.; Ibanez, B.; Group, E. S. C. S. D., 2023 ESC Guidelines for the management of acute coronary syndromes. Eur. Heart J. 2023, 44 (38), 3720–3826.

3. Backman, J. D.; O’Connell, J. R.; Tanner, K.; Peer, C. J.; Figg, W. D.; Spencer, S. D.; Mitchell, B. D.; Shuldiner, A. R.; Yerges-Armstrong, L. M.; Horenstein, R. B.; Lewis, J. P., Genome-wide analysis of clopidogrel active metabolite levels identifies novel variants that influence antiplatelet response. Pharmacogenet Genomics 2017, 27 (4), 159–163.

4. Jiang, X. L.; Samant, S.; Lesko, L. J.; Schmidt, S., Clinical pharmacokinetics and pharmacodynamics of clopidogrel. Clin. Pharmacokinet. 2015, 54 (2), 147–66.

5. Kazui, M.; Nishiya, Y.; Ishizuka, T.; Hagihara, K.; Farid, N. A.; Okazaki, O.; Ikeda, T.; Kurihara, A., Identification of the human cytochrome P450 enzymes involved in the two oxidative steps in the bioactivation of clopidogrel to its pharmacologically active metabolite. Drug metabolism and disposition 2010, 38 (1), 92–9.

6. Herbert, J. M.; Savi, P., P2Y12, a new platelet ADP receptor, target of clopidogrel. Semin Vasc Med 2003, 3 (2), 113–22.

7. Zhu, Y.; Zhou, J., In Vitro Biotransformation Studies of 2-Oxo-clopidogrel: Multiple Thiolactone Ring-Opening Pathways Further Attenuate Prodrug Activation. Chem Res Toxicol 2013, 26 (1), 179–190.

8. Angiolillo, D. J.; Jakubowski, J. A.; Ferreiro, J. L.; Tello-Montoliu, A.; Rollini, F.; Franchi, F.; Ueno, M.; Darlington, A.; Desai, B.; Moser, B. A.; Sugidachi, A.; Guzman, L. A.; Bass, T. A., Impaired responsiveness to the platelet P2Y12 receptor antagonist clopidogrel in patients with type 2 diabetes and coronary artery disease. J. Am. Coll. Cardiol. 2014, 64 (10), 1005–14.

9. Ancrenaz, V.; Daali, Y.; Fontana, P.; Besson, M.; Samer, C.; Dayer, P.; Desmeules, J., Impact of genetic polymorphisms and drug-drug interactions on clopidogrel and prasugrel response variability. Curr Drug Metab 2010, 11 (8), 667–77.

10. Wiviott, S. D.; Braunwald, E.; McCabe, C. H.; Montalescot, G.; Ruzyllo, W.; Gottlieb, S.; Neumann, F. J.; Ardissino, D.; De Servi, S.; Murphy, S. A.; Riesmeyer, J.; Weerakkody, G.; Gibson, C. M.; Antman, E. M.; Investigators, T.-T., Prasugrel versus clopidogrel in patients with acute coronary syndromes. N. Engl. J. Med. 2007, 357 (20), 2001–15.

11. Wallentin, L.; Becker, R. C.; Budaj, A.; Cannon, C. P.; Emanuelsson, H.; Held, C.; Horrow, J.; Husted, S.; James, S.; Katus, H.; Mahaffey, K. W.; Scirica, B. M.; Skene, A.; Steg, P. G.; Storey, R. F.; Harrington, R. A.; Investigators, P.; Freij, A.; Thorsen, M., Ticagrelor versus clopidogrel in patients with acute coronary syndromes. N. Engl. J. Med. 2009, 361 (11), 1045–57.

12. Kelly, R. P.; Close, S. L.; Farid, N. A.; Winters, K. J.; Shen, L.; Natanegara, F.; Jakubowski, J. A.; Ho, M.; Walker, J. R.; Small, D. S., Pharmacokinetics and pharmacodynamics following maintenance doses of prasugrel and clopidogrel in Chinese carriers of CYP2C19 variants. Br J Clin Pharmacol 2012, 73 (1), 93–105.

13. Lun, R.; Dhaliwal, S.; Zitikyte, G.; Roy, D. C.; Hutton, B.; Dowlatshahi, D., Comparison of Ticagrelor vs Clopidogrel in Addition to Aspirin in Patients With Minor Ischemic Stroke and Transient Ischemic Attack: A Network Meta-analysis. JAMA Neurol 2022, 79 (2), 141–148.

14. Pereillo, J. M.; Maftouh, M.; Andrieu, A.; Uzabiaga, M. F.; Fedeli, O.; Savi, P.; Pascal, M.; Herbert, J. M.; Maffrand, J. P.; Picard, C., Structure and stereochemistry of the active metabolite of clopidogrel. Drug Metab Dispos 2002, 30 (11), 1288–95.

15. Tuffal, G.; Roy, S.; Lavisse, M.; Brasseur, D.; Schofield, J.; Delesque Touchard, N.; Savi, P.; Bremond, N.; Rouchon, M. C.; Hurbin, F.; Sultan, E., An improved method for specific and quantitative determination of the clopidogrel active metabolite isomers in human plasma. Thromb. Haemost. 2011, 105 (4), 696–705.

16. Savi, P.; Pereillo, J. M.; Uzabiaga, M. F.; Combalbert, J.; Picard, C.; Maffrand, J. P.; Pascal, M.; Herbert, J. M., Identification and biological activity of the active metabolite of clopidogrel. Thromb. Haemost. 2000, 84 (5), 891–6.

17. Sun, Y.; Venugopal, J.; Guo, C.; Fan, Y.; Li, J.; Gong, Y.; Chen, Y. E.; Zhang, H.; Eitzman, D. T., Clopidogrel Resistance in a Murine Model of Diet-Induced Obesity Is Mediated by the Interleukin-1 Receptor and Overcome With DT-678. Arterioscler. Thromb. Vac. Biol. 2020, 40 (6), 1533–1542.

18. Zhang, H.; Lauver, D. A.; Wang, H.; Sun, D.; Hollenberg, P. F.; Chen, Y. E.; Osawa, Y.; Eitzman, D. T., Significant Improvement of Antithrombotic Responses to Clopidogrel by Use of a Novel Conjugate as Revealed in an Arterial Model of Thrombosis. J Pharmacol Exp Ther 2016, 359 (1), 11–7.

19. Lauver, D. A.; Kuszynski, D. S.; Christian, B. D.; Bernard, M. P.; Teuber, J. P.; Markham, B. E.; Chen, Y. E.; Zhang, H., DT-678 inhibits platelet activation with lower tendency for bleeding compared to existing P2Y12 antagonists. Pharmacol Res Perspect 2019, 7 (4), e00509.

20. Liu, Z.; Liu, S.; Gong, Y.; Chi, X.; Wang, T.; Fan, F.; Qu, C.; Lou, Y.; Zhang, L.; Zhang, B.; Yang, F.; Mohetaboer, M.; Wang, J.; Qiu, L.; Miao, L.; Lu, Y.; You, R.; He, P.; Li, Y.; Yi, T.; Weng, H.; Xia, Y.; Wang, C.; Shi, Q.; Wang, Z.; Jiang, Y.; Li, Y.; Han, C.; Wang, Y.; Wang, X.; Yang, C.; Chen, Y. E.; Eitzman, D. T.; Zhang, H.; Li, J., The first in-human study to evaluate the antiplatelet properties of the clopidogrel conjugate DT-678 in acute coronary syndrome patients and healthy volunteers. Br J Pharmacol 2025, 182 (1), 131–141.

21. Shaw, S. A.; Balasubramanian, B.; Bonacorsi, S.; Cortes, J. C.; Cao, K.; Chen, B. C.; Dai, J.; Decicco, C.; Goswami, A.; Guo, Z.; Hanson, R.; Humphreys, W. G.; Lam, P. Y.; Li, W.; Mathur, A.; Maxwell, B. D.; Michaudel, Q.; Peng, L.; Pudzianowski, A.; Qiu, F.; Su, S.; Sun, D.; Tymiak, A. A.; Vokits, B. P.; Wang, B.; Wexler, R.; Wu, D. R.; Zhang, Y.; Zhao, R.; Baran, P. S., Synthesis of Biologically Active Piperidine Metabolites of Clopidogrel: Determination of Structure and Analyte Development. J Org Chem 2015, 80 (14), 7019–32.

22. Bluet, G.; Blankenstaein, J.; Brohan, E.; Prevost, C.; Cheve, M.; Schofield, J.; Roy, S., Synthesis of the stablized active metabolite of clopidogrel. Tetrahedron Lett. 2014, 70, 3893–3900.

23. Huang, X.; Zhou, J.; Yang, D.; Zhang, J.; Xia, X.; Chen, Y. E.; Xu, J., Decoding CRISPR-Cas PAM recognition with UniDesign. Brief Bioinform 2023, 24 (3).

24. Huang, X.; Sun, Y.; Osawa, Y.; Chen, Y. E.; Zhang, H., Computational redesign of cytochrome P450 CYP102A1 for highly stereoselective omeprazole hydroxylation by UniDesign. J Biol Chem 2023, 299 (8), 105050.

25. Butler, C. F.; Peet, C.; Mason, A. E.; Voice, M. W.; Leys, D.; Munro, A. W., Key mutations alter the cytochrome P450 BM3 conformational landscape and remove inherent substrate bias. J Biol Chem 2013, 288 (35), 25387–99.

26. Bendich, A.; Machlin, L. J.; Burton, G. W.; Wayner, D. D. M., The antioxidant role of vitamin C. Advances in Free Radical Biology & Medicine 1986, 2 (2), 419–444.

27. Fabelle, N. R.; Oktavia, F.; Cha, G. S.; Nguyen, N. A.; Choi, S. K.; Yun, C. H., Production of a major metabolite of niclosamide using bacterial cytochrome P450 enzymes. Enzyme Microb. Technol. 2023, 165, 110210.

28. Harkey, A.; Kim, H. J.; Kandagatla, S.; Raner, G. M., Defluorination of 4-fluorophenol by cytochrome P450(BM(3))-F87G: activation by long chain fatty aldehydes. Biotechnology letters 2012, 34 (9), 1725–31.

29. Pettersen, E. F.; Goddard, T. D.; Huang, C. C.; Couch, G. S.; Greenblatt, D. M.; Meng, E. C.; Ferrin, T. E., UCSF Chimera--a visualization system for exploratory research and analysis. J Comput Chem 2004, 25 (13), 1605–12.

30. Huang, X.; Pearce, R.; Zhang, Y., EvoEF2: accurate and fast energy function for computational protein design. Bioinformatics 2020, 36 (4), 1135–1142.

31. Shapovalov, M. V.; Dunbrack, R. L., Jr., A smoothed backbone-dependent rotamer library for proteins derived from adaptive kernel density estimates and regressions. Structure 2011, 19 (6), 844–58.

32. Omura, T.; Sato, R., The carbon monoxide-binding pigment of liver microsomes. I. Evidence for Its hemoprotein nature. J Biol Chem 1964, 239, 2370–8.

33. Gay, C.; Collins, J.; Gebicki, J. M., Determination of hydroperoxides by the ferric-xylenol orange method. Redox Rep 1999, 4 (6), 327–8.

34. Kuthan, H.; Ullrich, V.; Estabrook, R. W., A quantitative test for superoxide radicals produced in biological systems. Biochem J 1982, 203 (3), 551–8.

35. Gruenke, L. D.; Konopka, K.; Cadieu, M.; Waskell, L., The stoichiometry of the cytochrome P-450-catalyzed metabolism of methoxyflurane and benzphetamine in the presence and absence of cytochrome b5. J Biol Chem 1995, 270 (42), 24707–18.

